# Neurofluid Coupling during Sleep and Wake States

**DOI:** 10.1101/2022.10.31.514639

**Authors:** Vidhya Vijayakrishnan Nair, Brianna R Kish, Pearlynne L H Chong, Ho-Ching (Shawn) Yang, Yu-Chien Wu, Yunjie Tong, A. J. Schwichtenberg

## Abstract

Low-frequency changes in cerebral hemodynamics have recently been shown to drive cerebrospinal fluid (CSF) movement in the human brain during non-rapid eye movement (NREM) sleep and resting state wakefulness. However, whether the coupling strength between these neurofluids varies between wake and sleep states is not known. In addition, the principal origin (i.e., neuronal vs. systemic) of these slow cerebral hemodynamic oscillations in either state also remains unexplored. To investigate this, a wake/sleep study was conducted on eight young, healthy volunteers, concurrently acquiring neurofluid dynamics using functional Magnetic Resonance Imaging, neural activity using Electroencephalography, and non-neuronal systemic physiology with peripheral functional Near-Infrared Spectroscopy. Our results reveal that low-frequency cerebral hemodynamics and CSF movements are strongly coupled regardless of whether participants were awake or in light NREM sleep. Furthermore, it was also found that, while autonomic neural contributions are present only during light NREM sleep, non-neuronal systemic physiology influences neurofluid low-frquency oscillations in a significant way across both wake and sleep states. These results further our understanding regarding the low-frequency hemodynamic drivers of CSF movement in the human brain and could help inform the development of therapies for enhancing CSF circulation.

## 1. Introduction

Cerebrospinal fluid (CSF) movement within the central nervous system (CNS) is critical for optimal brain health and function. CSF movement aids brain homeostasis by transporting nutrients, hormones, and other immune system components through the CNS and provides mechanical protection and support to the brain and spinal cord^1–3^. In addition, several recent studies on CSF movement highlight its role in the pathophysiology of neurodegenerative^4,5^ and neurodevelopmental^6–8^ disorders and the glymphatic system^9^.

Current theories on the form and functions of the glymphatic system posit that CSF movement/circulation is crucial for disseminating growth factors and removing metabolic wastes from the brain ^10^. With increased CSF production and circulation during sleep, CSF dynamics are likely linked with sleep processes, especially slow wave NREM sleep^11^. For example, Fultz *et al*., in a functional Magnetic Resonance Imaging (fMRI) study, reported local electrocortical slow wave activity (SWA) initiated control of low-frequency (LF) cerebral hemodynamic changes and consequent large CSF movement during NREM sleep (in humans). This study reported a sequential occurrence of neural SWA peaks shortly followed by LF cerebral blood volume decreases and large cranially directed CSF movement at the fourth ventricle, exclusively during NREM sleep^12^. Moreover, this study reported significant increases in CSF movement variation during NREM sleep compared to resting state wakefulness.

In another human fMRI study, Yang *et al*., observed large CSF movement in both directions (i.e., craniad and caudad) coupled with cerebral blood volume changes in the LF range (during resting state wakefulness). Yang *et al*. posited a mechanical coupling model between these neurofluids1 based on the Monro-Kellie doctrine. However, this study did not explore the neural or physiological control behind the documented neurofluid coupling. Given these observations, it is unclear whether the coupling strength between these neurofluids in the LF range significantly differs between resting state wakefulness and NREM sleep.

A recent human fMRI study by Picchioni *et al*. reported a similar sleep-related SWA control of cranially directed CSF movement and explored the driving forces behind these effects. Results suggest an autonomic rather than local cortical control of coupled neurofluid oscillations, particularly during light NREM sleep^13^. In addition to coupled changes in neurofluids, they also reported associated changes in autonomic physiology (visible via transient changes in heart activity and respiration) in response to autonomic arousals during light NREM sleep. Notably, global systemic circulatory sources (independent of neural control) are also documented contributors to LF cerebrovascular oscillations. Examples include variations in arterial partial pressure of CO_2_^14,15^ and spontaneous vasomotion^16,17^. These diverse observations of LF ‘drivers’ warrant further investigation; specifically in the area of neurofluid coupling during resting state wakefulness and NREM sleep – i.e., whether neurofluid coupling reflects more neuronal or systemic physiological contributions and more importantly, do these contributions change between resting state wakefulness and light NREM sleep.

In the present study, we intend to answer the following questions (see Figure 1 for a pictorial summary): (1) Does CSF movement variation (indexed at the fourth ventricle) increase significantly during light NREM sleep when compared to resting wakefulness? (2) Does the coupling strength between neurofluids change significantly during light NREM sleep (compared to resting wakefulness)? (3) What is the principal cause or driving force behind this neurofluid coupling across states? To answer this question, we explored neurofluid movement with a neurogenic origin (through the relationship between neurofluids and electrical activity of the brain) and/or a global circulatory origin (through the relationship between neurofluids and non-neuronal fingertip vascular LFOs).

**Figure 1:**
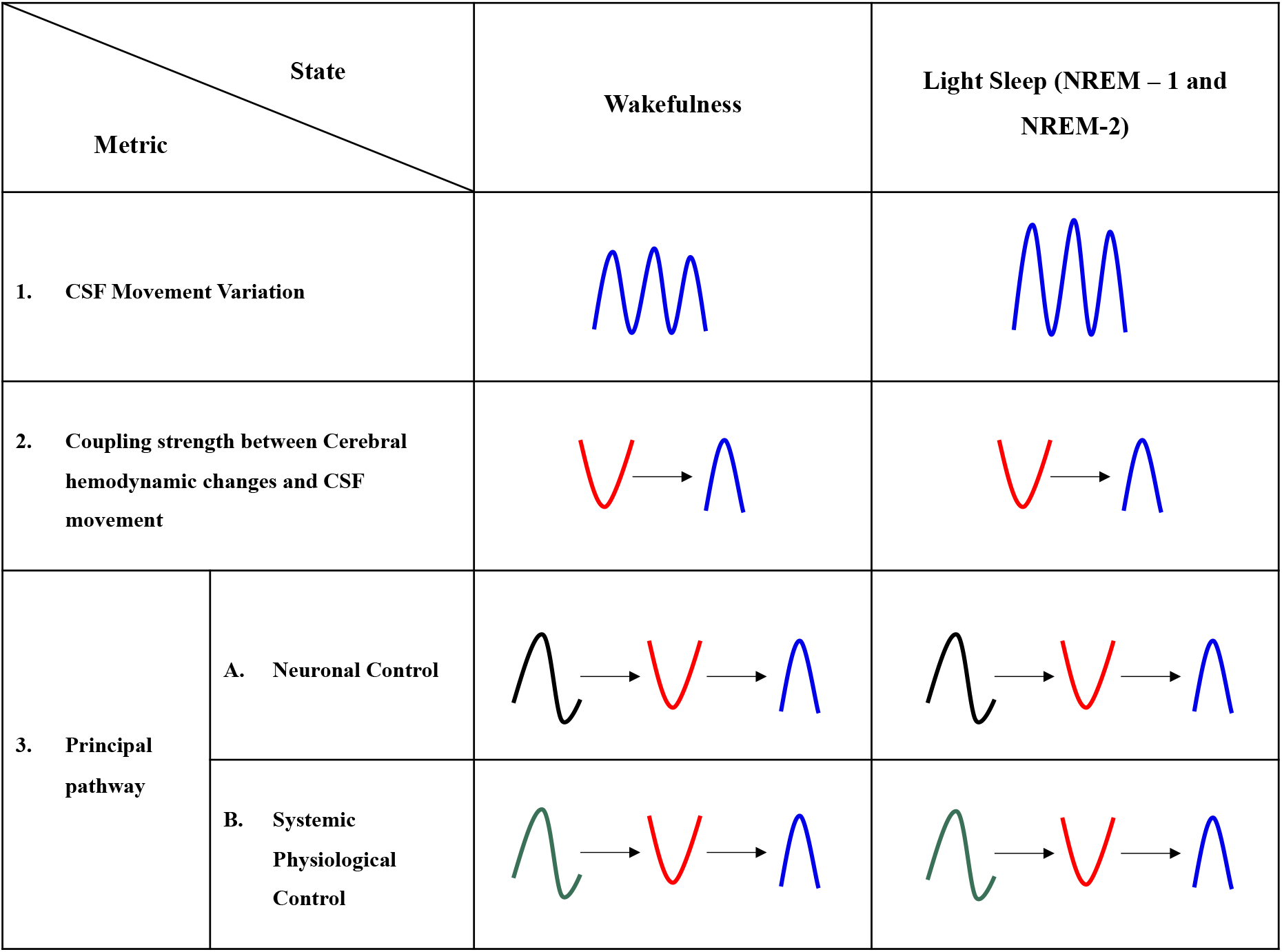
Summary of research questions. 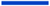 CSF Movement 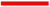 Cerebral hemodynamic Changes 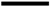 EEG 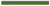 Non-neuronal systemic physiology; CSF - Cerebrospinal Fluid; EEG – Electroencephalogram; NREM – Non Rapid Eye Movement sleep.

It is critical to understand neurofluid coupling across wake/sleep states for two reasons: First, if neurofluid coupling is consistent across sleep and wake states, then future studies can use the least intrusive metric to estimate CSF movement. Second, if there is a shift in CSF movement drivers across sleep and wake states, then therapeutics designed to improve CSF circulation will have neuronal or physiological/circulatory targets. To address our research questions, we designed and implemented a wake/sleep study with simultaneous acquisition of neurofluid dynamics with fMRI, electrical brain activity with Electroencephalography (EEG), and non-neuronal systemic physiology with peripheral (fingertip) functional Near-Infrared Spectroscopy (fNIRS).

## 2. Materials and Methods

### 2.1. Participants

Eight healthy participants (4 females and 4 males) aged 20 – 25 (21.4 ± 1.7) years were included in this study. The study was approved by Purdue University’s Human Research Protection Plan (IRB-2020-1329) and was conducted in accordance with the application of Belmont Report principles (Respect for Persons, Beneficence, and Justice) and federal regulations 45 CFR 46, 21 CFR 50, 56. Written informed consent was obtained from all participants.

### 2.2. Experimental Design

A pictorial description of the experimental design and data analysis methods is illustrated in Figure 2.

**Figure 2:**
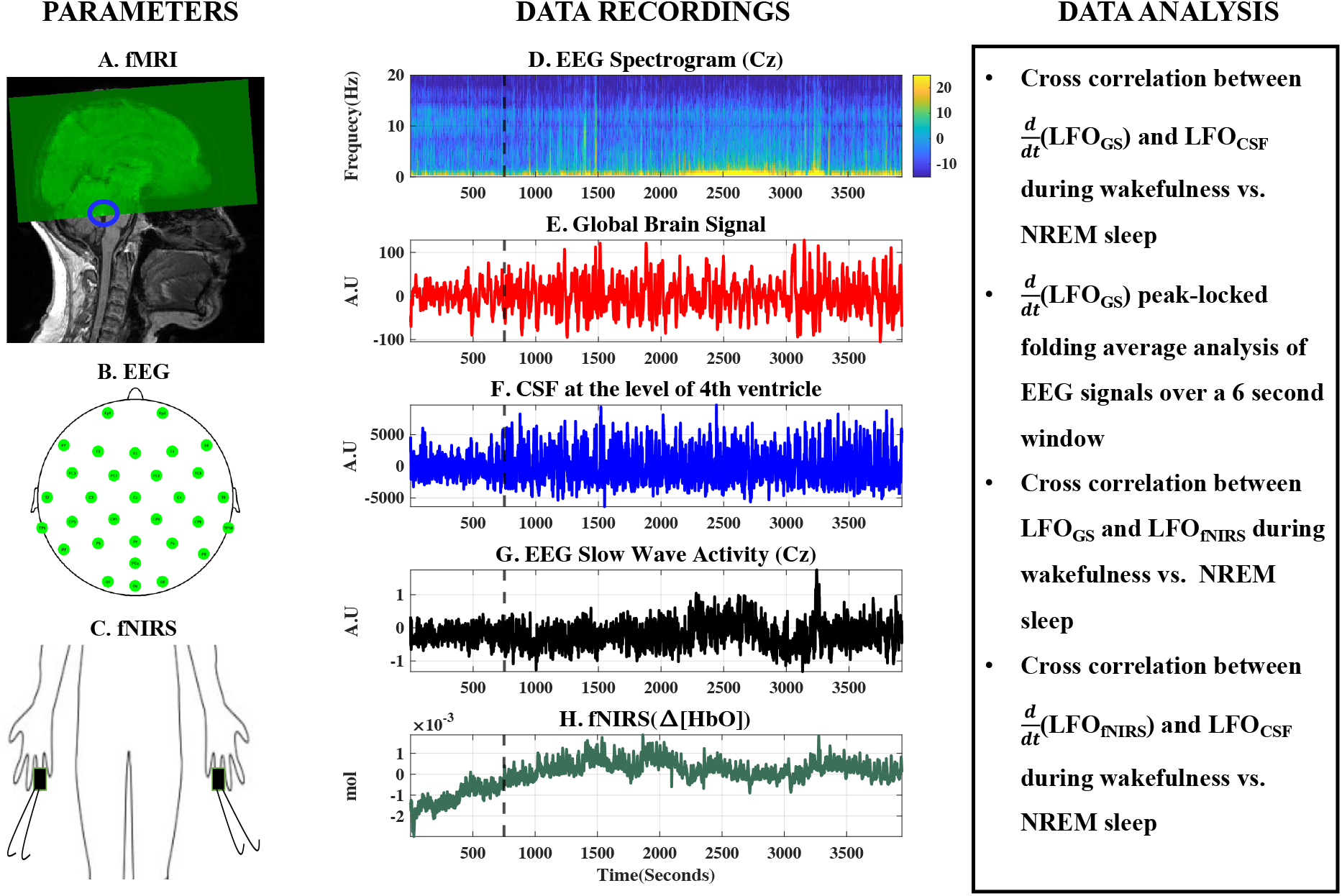
Experimental Design and Data Analysis Stream. The considered parameters, data recordings and data analysis techniques employed to study the coupling between neurofluid oscillations are briefed in the first, second and third columns respectively. (A) Typical fMRI scan design illustrating the capture of cranially directed CSF movement into the brain scan volume utilizing inflow effect and the corresponding raw (E) global brain signal and (F) CSF movement signal. (B) Locations of 32-channel EEG electrodes on the head as well as (D) EEG spectrogram and (G) Slow wave Activity from the channel Cz. (C) Placement of fingertip fNIRS sensors and (H) raw fNIRS (Δ[HbO]) signal. The black dashed vertical line represents the starting point of NREM sleep. A.U – Arbitrary units; CSF - Cerebrospinal Fluid; EEG – Electroencephalogram; fMRI – functional magnetic resonance imaging; fNIRS – functional near infrared spectroscopy; Δ[HbO] – Changes in concentration of oxyhemoglobin; LFO – Low frequency oscillations (0.01 Hz – 0.1 Hz); NREM – Non Rapid Eye Movement sleep.

#### 2.2.1. Concurrent EEG, MRI, and fNIRS Data Acquisition

All participants had regular habitual sleep schedules (confirmed with a minimum of four 24-hour sleep/wake cycles via actigraphy). The data collection/experimental session was scheduled for 4 hours past their typical nighttime sleep onset time. During the experimental session, the participants wore an MRI-compatible EEG cap (Braincap MR, Brain Products GmbH, Gilching, Germany) and were requested to sleep inside a 3T SIEMENS MRI scanner (Magnetom Prisma, Siemens Medical Solutions, Erlangen, Germany) with a 64-channel head coil. fMRI scans (Figure 2A) of the brain were acquired using the following parameters: FOV = 230 mm, acquisition matrix = 92 × 92, 48 slices, voxel size = 2.5 × 2.5 × 2.5 mm, TR/TE = 960/30.6 ms, echo-spacing = 0.51 ms, flip angle = 35°, multiband acceleration factor = 8, multi-slice mode: interleaved. TR was changed to 480 ms in two of the participants. Concurrent 32-channel EEG data (Figure 2B) was recorded using the MRI-compatible EEG system (BrainAmp MR, Brain Products GmbH, Gilching, Germany) at a sampling rate of 5000 Hz, with FCz electrode as the reference. The dropdown ECG electrode from the EEG cap was attached paravertebrally on the left back of the participants. The skin-electrode impedance was maintained below 10 KΩ during the recording for all electrodes. In addition, concurrent peripheral hemodynamics were recorded (Figure 2C) from two fingertips (fourth finger of both arms) using a Continuous Wave (CW) fNIRS system (NIRScoutXP NIRx Medizintechnik GmbH, Berlin, Germany) with two laser sources, each combining two wavelengths (785 and 830 nm) and MRI-compatible fNIRS probes with 10 m long optical fibers. All participants also wore a chest belt to measure respiration.

The fMRI scans were designed to capture the cranially directed CSF movement utilizing the inflow effect^12^. As illustrated in Figure 2A, by placing the bottom edge of the scan volume appropriately at the fourth ventricle, an intense CSF flow signal directed into the scan volume can be obtained. Consequently, the slice of interest at the fourth ventricle was always acquired first in the scan. We chose to evaluate the CSF movement at the level of the fourth ventricle because the narrow, tapered structure limits the movement of CSF in other directions, thereby strengthening the inflow effect. Prior to sleep onset within the scanner, structural T1-weighted MPRAGE (Magnetization Prepared Rapid Acquisition Gradient Echo) images (TR/TE: 2300/2.26 ms, 192 slices per slab, flip angle: 8°, resolution: 1.0mm × 1.0mm × 1.0mm) and structural 3D SPACE (Sampling Perfection with Application optimized Contrasts using different flip angle Evolution) T2-weighted images (TR/TE: 2800/409 ms, 208 slices per slab, resolution: 0.8mm × 0.8mm × 0.7mm) were also acquired.

#### 2.2.2. Data Preparation

##### EEG preprocessing and sleep scoring

EEG signals were initially cleaned by removing MR gradient artifacts and cardioballistic artifacts using the average artifact subtraction method^18^ and downsampled to 250 Hz using Analyzer software (Brain Vision, Morrisville, USA). Independent Component Analysis was also performed afterward to remove the ocular, muscular, and remaining electrocardiographic artifacts using EEGLAB v2021.0 toolbox^19^. Further, sleep stages (NREM-1, NREM-2, NREM-3, REM) were scored in 30-second epochs using the American Academy of Sleep Medicine scoring manual criteria^20^. Ample amounts of wakefulness, NREM-1, and NREM-2 were captured and utilized for this study. NREM-3 and REM were not captured consistently across participants, and only small segments were available for analysis; therefore, the present study focuses on wakefulness, NREM-1, and NREM-2.

##### fMRI preprocessing and signal extraction

fMRI data were processed using FSL (FMRIB Expert Analysis Tool, v6.01; Oxford University, UK^21^) and MATLAB (MATLAB 2021b, The MathWorks Inc., Natick, MA, 2000). First, the CSF signal was extracted from a voxel in the fourth ventricle identified by overlaying the fMRI data over the structural T1-weighted image registered to the fMRI data (see Figure 2F). Care was taken to extract the signal from the center of the fourth ventricle so that it comes from a voxel with negligible partial-volume effects from surrounding tissues. The fMRI data were only corrected for slice-timing (FSL SLICETIMER) before extracting the single voxel CSF signal. No motion correction was applied^12,22^. To confirm that the CSF signals were not corrupted by motion, we assessed and documented no significant correlations between motion parameters (FSL MCFLIRT) and the CSF signals (see table S1 in the supplementary material). Having obtained the CSF movement signal from the fourth ventricle without motion correction, motion correction (FSL MCFLIRT) and slice-timing correction (FSL SLICETIMER) were applied to fMRI data used in subsequent analyses. Further, the global mean of fMRI signal (GS) was extracted across the whole brain (Figure 2E).

#### 2.2.3. Data Analysis

##### Research Question 1-CSF movement during wakefulness and light NREM sleep

To capture sleep-dependent CSF movement fluctuations, the standard deviation (SD) of 5-minute segments was quantified for each wake/sleep stage. First, one-sample Kolmogorov-Smirnov test was used to test for normality in each wake/sleep stage. Second, non-parametric Kruskal-Wallis tests were used to assess SD differences across the wake/sleep stages.

##### Research Question 2 – Neurofluid coupling during wakefulness and light NREM sleep

CSF movement signals were linearly detrended, demeaned, and bandpass filtered to the range of 0.01 Hz – 0.1 Hz, using a zero delay fourth-order Butterworth filter to extract the corresponding low-frequency oscillations (LFOs). As we demonstrated in our previous work^22^, the changes in cerebral blood volume (CBV) rather than CBV itself are a driving force of CSF movement. Without the CBV changes, no force will be exerted on the ventricles. Therefore, instead of the fMRI GS signal itself, we use the derivative of the GS to reflect CBV changes in the brain. Further, maximum cross-correlation coefficients (MCCCs) and corresponding time delays for each participant were calculated (MATLAB xcorr, maximum lag range: ±15 seconds) between 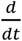 (LFO_GS_) and LFO_CSF_ (calculations were performed across continuous 5-minute segments of each stage). The absolute maximum value from the calculated CCCs was identified as the MCCC with its original arithmetic sign. Based on previous research, only the MCCCs above the statistically established threshold of 0.3 (p-value<0.01 for positive MCCCs) or below -0.3 (p-value<0.01 for negative MCCCs) are considered significant in the LFO range^23,24^. MCCCs were not normally distributed (determined via one-sample Kolmogorov-Smirnov tests). Therefore, a non-parametric one-sample Wilcoxon signed rank test against 0.3/-0.3 were applied to MCCCs in the LFO range.

##### Research Question 3A – Local neuronal control assessed via EEG folding average analysis

This analysis was employed to detect the presence of neural activity prior to LF CSF movement in both directions during light NREM sleep. The Cz electrode was chosen to evaluate the SWA during NREM sleep stages because it has comparatively lower amplitude variability in the SWA range during NREM sleep. In addition, the location of the Cz electrode makes it less susceptible to ballistocardiogram artifacts caused by motion in the magnetic field, as it is held firmly against the MRI head holder. EEG SWA during NREM sleep was calculated by following the method described by Fultz *et al*. Briefly, the EEG signals were bandpass filtered in the range of 0.2 Hz – 4 Hz (eegfilt.m, EEGLAB v2021.0 toolbox), and the instantaneous amplitude envelope of the filtered signals (absolute value of the Hilbert transform) was extracted. Finally, the instantaneous amplitude time series were smoothed with a moving average of 4 seconds^12^ (see Figure 1G). A peak locked folding average analysis based on 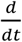 (LFO_GS_) peaks were employed here to identify EEG SWA prior to CSF inward/outward movement. Our previous research has illustrated that the negative and positive half of 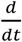 (LFO_GS_) can be used as a surrogate for inward (cranially directed) and outward (caudally directed) CSF movements, respectively at the level of the 4th ventricle^22,25^. Therefore, in this study, local maxima of 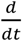 (LFO_GS_) signal that surpassed a threshold of 50% amplitude in the negative and positive direction are used as surrogates for CSF inward (cranially directed) and outward (caudally directed) movements at the level of the 4^th^ ventricle respectively. Further, the EEG SWA for a duration of 6 seconds prior to each of the identified local maxima in the 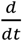 (LFO_GS_) signal was averaged. This averaging was performed separately in both directions to detect EEG SWA in both movement directions. As a comparison, a similar peak locked folding average analysis was also performed during wakefulness by using the full-spectrum EEG signals (bandpass-filtered in the range of 0.1 Hz – 45 Hz) from the same electrode (Cz).

##### Research Question 3B – Systemic physiological control assessed via fNIRS signal and neurofluid coupling

The fNIRS change in concentration of oxyhemoglobin (Δ[HbO]) signals, GS and CSF movement signals were linearly detrended, demeaned and bandpass filtered to the range of 0.01 Hz – 0.1 Hz, using a zero delay fourth-order Butterworth filter to extract the corresponding LFOs. The peripheral hemodynamic LFOs from both fingertips exhibited high correlations (0.72±0.23). Hence, these signals were averaged for each participant to increase the signal-to-noise ratio (SNR). Further, MCCCs and corresponding time delays for each participant were calculated (MATLAB xcorr, maximum lag range: ±15 seconds) between 1. LFO_fNIRS_ and LFO_GS_ and 2. 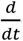 (LFO_fNIRS_) and LFO_CSF_ (calculations were performed across continuous 5-minute segments of each stage). MCCC was identified as the absolute maximum value from the calculated CCCs, with its original arithmetic sign. As discussed before, a non-parametric one-sample Wilcoxon signed rank test against the established statistical threshold of 0.3/-0.3 was applied on MCCCs in the LFO range since they were not normally distributed (one sample Kolmogorov-Smirnov test was used to test for normality).

## 3. Results

### 3.1. Research Question 1 - Does CSF movement variation increase during light NREM sleep?

As illustrated in Figure 3, there was only a modest increase in CSF movement during light NREM sleep compared to wakefulness. Precisely, CSF movement into the brain from the spinal canal captured at the level of the fourth ventricle only had a moderate increase from resting state wakefulness to NREM 1 and NREM 2. Figure 3A illustrates a 5-minute signal segment of each stage for a representative participant. The flow-weighted CSF movement peaks in the cranial direction were clearly present across all the stages. The group results (Figure 3B) of standard deviations of continuous 5-minute signal segments from all participants illustrated no significant differences across wakefulness, NREM-1, and NREM-2.

**Figure 3:**
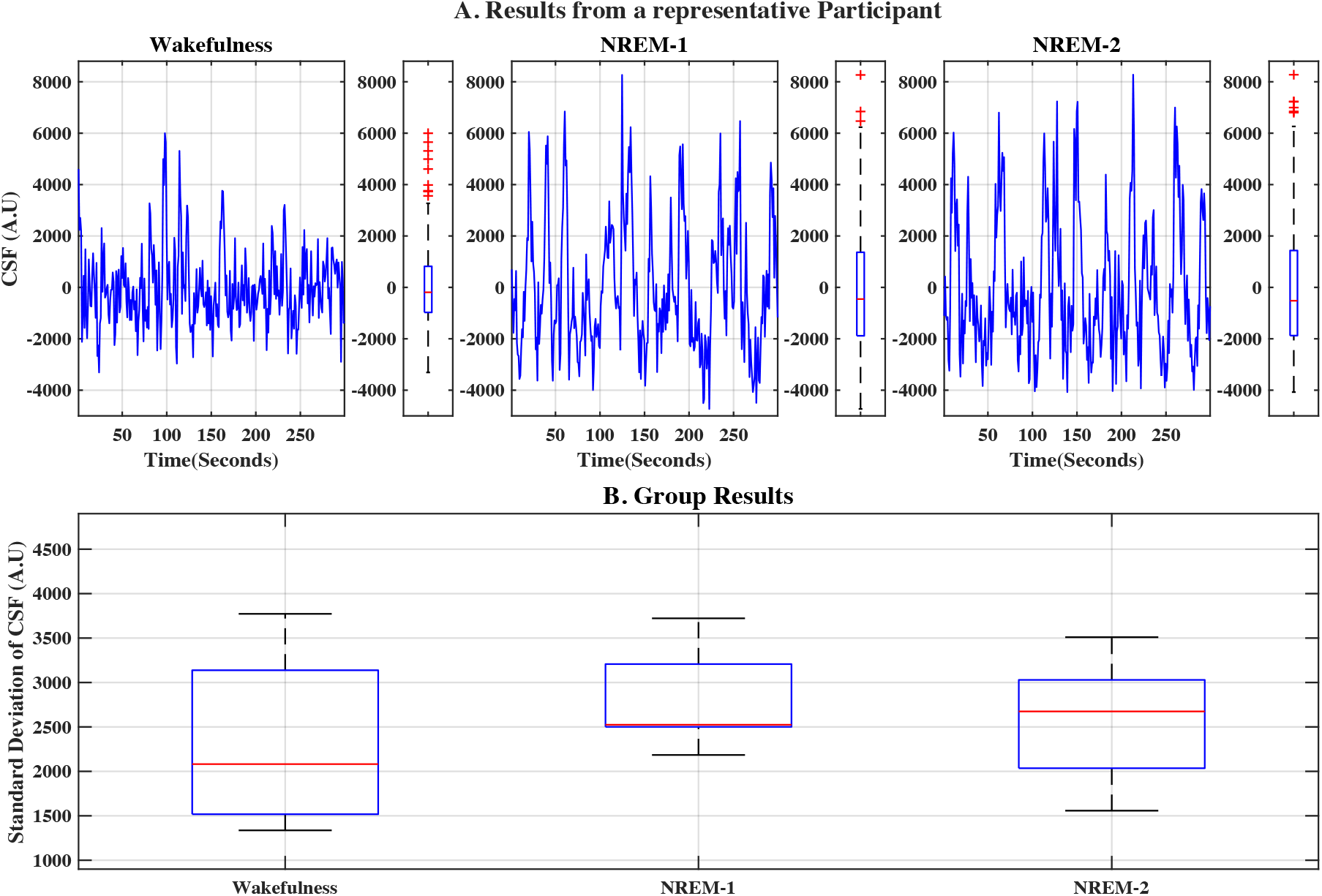
Changes in cranially directed CSF movement variation between wakefulness vs. NREM-1 and NREM-2. (A) An example of continuous 5-minute segments of CSF movement and their distribution across all three stages from a representative participant and (B) group results of standard deviations of continuous 5-minute signal segments from all participants. A.U – Arbitrary Units; CSF - Cerebrospinal Fluid; NREM – Non-Rapid Eye Movement sleep

### 3.2. Research Question 2 – Does the coupling strength between neurofluids change during light NREM sleep?

No significant changes in coupling strength between LF brain hemodynamic changes and CSF movement were documented during light NREM sleep when compared to wakefulness. The results of the cross-correlation analysis between LF brain hemodynamic changes and CSF movement in the LFO range across all three stages are illustrated in Figure 4.

**Figure 4:**
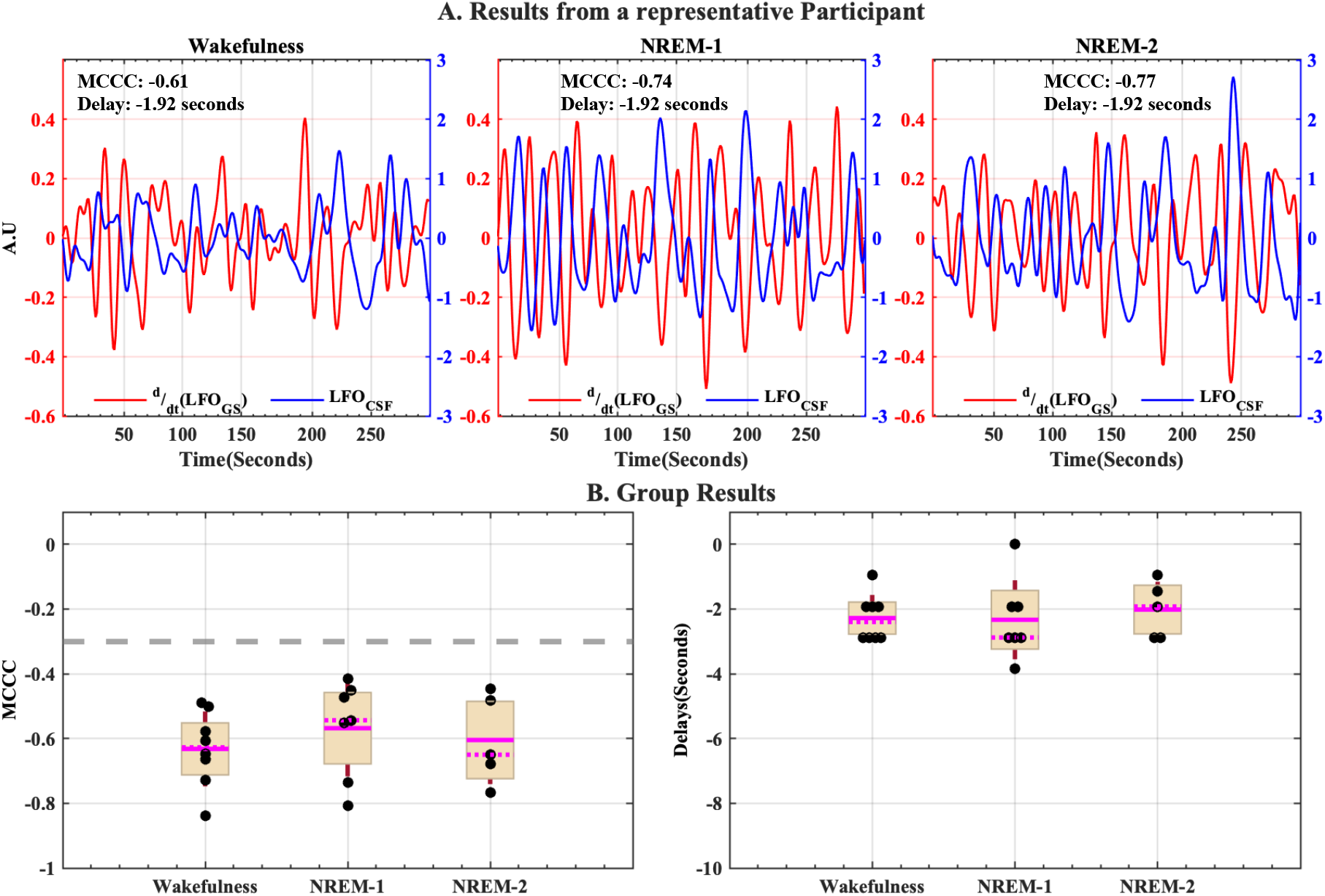
Strong coupling exists between LF brain hemodynamic changes and cranially directed CSF movement across all three stages. Results of MCCCs and corresponding delays between 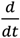 (LFO_GS_) and LFO_CSF_ during continuous 5-minute segments of wakefulness, NREM-1 and NREM-2 for (A) a representative participant and (B) for all participants enrolled in the study. A.U – Arbitrary Units; CSF - Cerebrospinal Fluid; GS – Global Signal; LFO – Low frequency Oscillation (0.01 Hz – 0.1 Hz); MCCC – Maximum Cross-Correlation Coefficient; NREM – Non-Rapid Eye Movement sleep. In group results (B), the magenta solid line represents the mean, the magenta dotted line represents the median, the brown whiskers represent one standard deviation of the raw data points jittered over a 95 percent confidence interval in cream and the gray dashed line represent the threshold of statistical significance for MCCCs in the LFO range.

MCCCs and corresponding delay times between 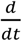 (LFO_GS_) and LFO_CSF_ during continuous 5-minute segments of wakefulness, NREM-1 and NREM-2 from a representative participant are represented in Figure 4A.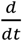 (LFO_GS_) exhibit significant negative correlations (MCCC < -0.3) with LFO_CSF_, at a time delay of -1.92 seconds during wakefulness, NREM-1, and NREM-2. The negative delay indicates that the 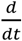 (LFO_GS_) signal leads the LFO_CSF_. Across all participants (Figure 4B), no significant changes were observed in mean negative correlations (wakefulness: -0.63±0.12; NREM-1: -0.57±0.15; NREM-2: -0.61±0.14) and in time delays (wakefulness: -2.28±0.71 seconds; NREM-1: -2.33±1.22 seconds; NREM-2: -2.02±0.86 seconds) between wakefulness, NREM-1 and NREM-2.

### 3.3. Research Question 3A – What is the principal cause or predominate driving force behind this neurofluid coupling in each assessed wake and sleep state? (Neuronal origin analyses)

EEG SWA occurs prior to LF brain hemodynamic changes in both directions during NREM sleep but not during wakefulness. The results of 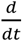 (LFO_GS_) peak-locked folding average analysis of EEG signals is illustrated in Figure 5. Clear EEG SWA peaks occurred for about 2 – 4 seconds and 2 – 3 seconds, respectively, before peaks in 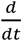 (LFO_GS_) in both directions (i.e., positive change of 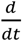 (LFO_GS_) before cranially directed inward CSF movement and negative change of 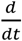 (LFO_GS_) before caudally directed outward CSF movement) during NREM-1 and NREM-2. However, no clear neural activity peaks (full spectrum EEG (Cz): 0.1 Hz – 45 Hz) were detected for a duration of 6 seconds before 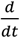 (LFO_GS_) peaks in either direction during resting state wakefulness.

**Figure 5:**
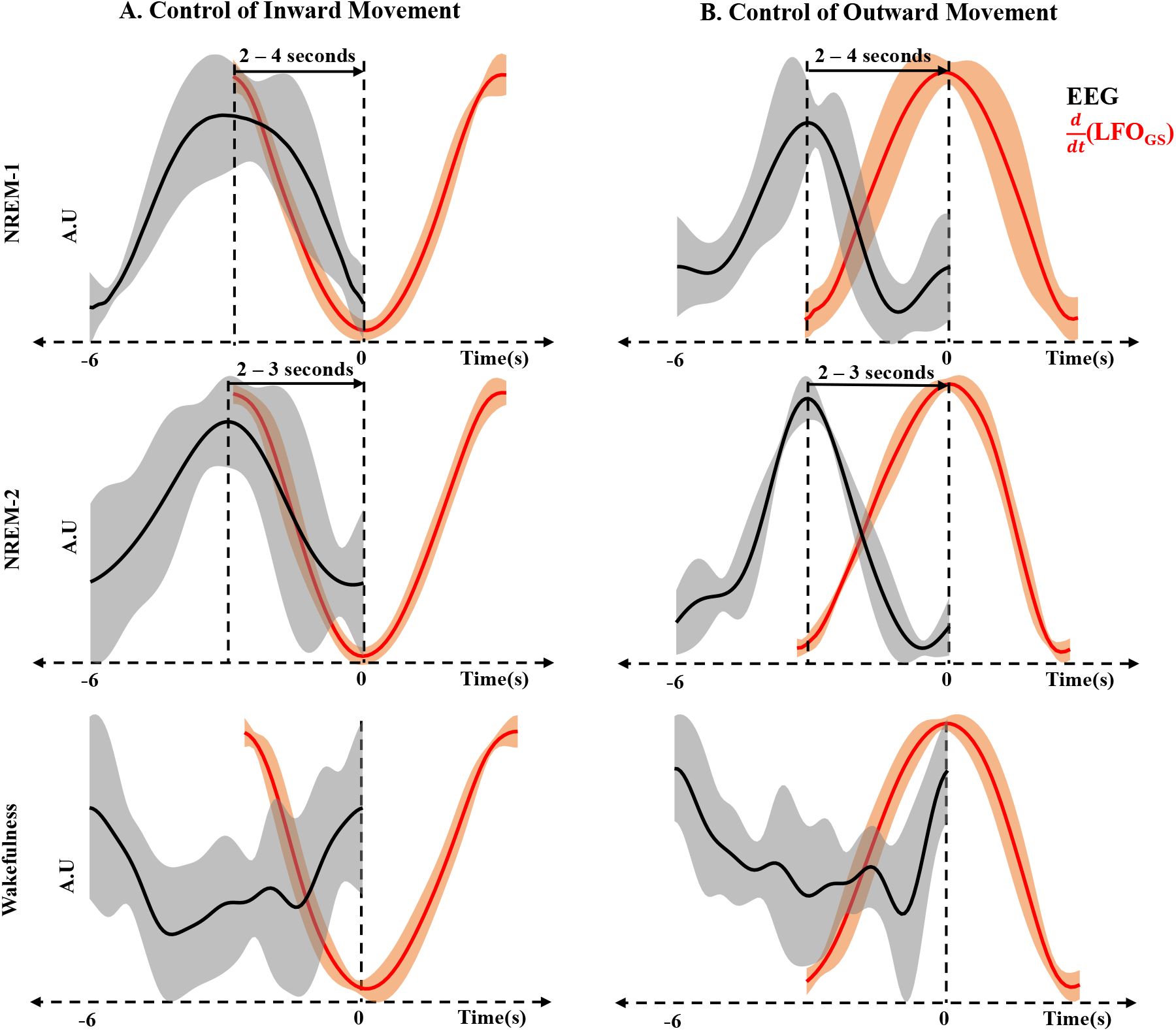
Results of 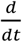 peak-locked folding average analysis of EEG signals in (A) positive direction (cranially directed CSF inward movement) and (B) negative direction (caudally directed outward CSF movement) during NREM-1, NREM-2 and wakefulness. The mean signal in each case is illustrated with thick black and red lines with standard deviation across participants represented by gray and orange regions around the mean signal. A.U – Arbitrary Units; CSF - Cerebrospinal Fluid; EEG – Electroencephalogram; GS – Global Signal; LFO – Low frequency Oscillation (0.01 Hz – 0.1 Hz); NREM – Non Rapid Eye Movement sleep.

### 3.4. Research Question 3B – What is the principal cause or predominate driving force behind neurofluid coupling in each assessed wake and sleep state? (Systemic circulatory origin analyses)

The results of the cross-correlation analysis between LF peripheral and brain hemodynamics and between LF peripheral hemodynamic changes and cranially directed CSF movement across all three stages are illustrated in Figure 6. MCCCs and corresponding delay times between LFO_fNIRS_ and LFO_GS_ during continuous 5-minute segments of wakefulness, NREM-1, and NREM-2 from a representative participant are presented in Figure 6A. LFO_fNIRS_ exhibited strong positive correlations (MCCC > 0.3) with LFO_GS_ during wakefulness, NREM-1, and NREM-2. Across all participants (Figure 6B), similar significant mean positive correlations (wakefulness: 0.61±0.16; NREM-1: 0.66±0.18; NREM-2: 0.62±0.14) were seen during wakefulness, NREM-1, and NREM-2. Time delays associated with the strong positive MCCCs are inconsistent across participants in all stages.

**Figure 6:**
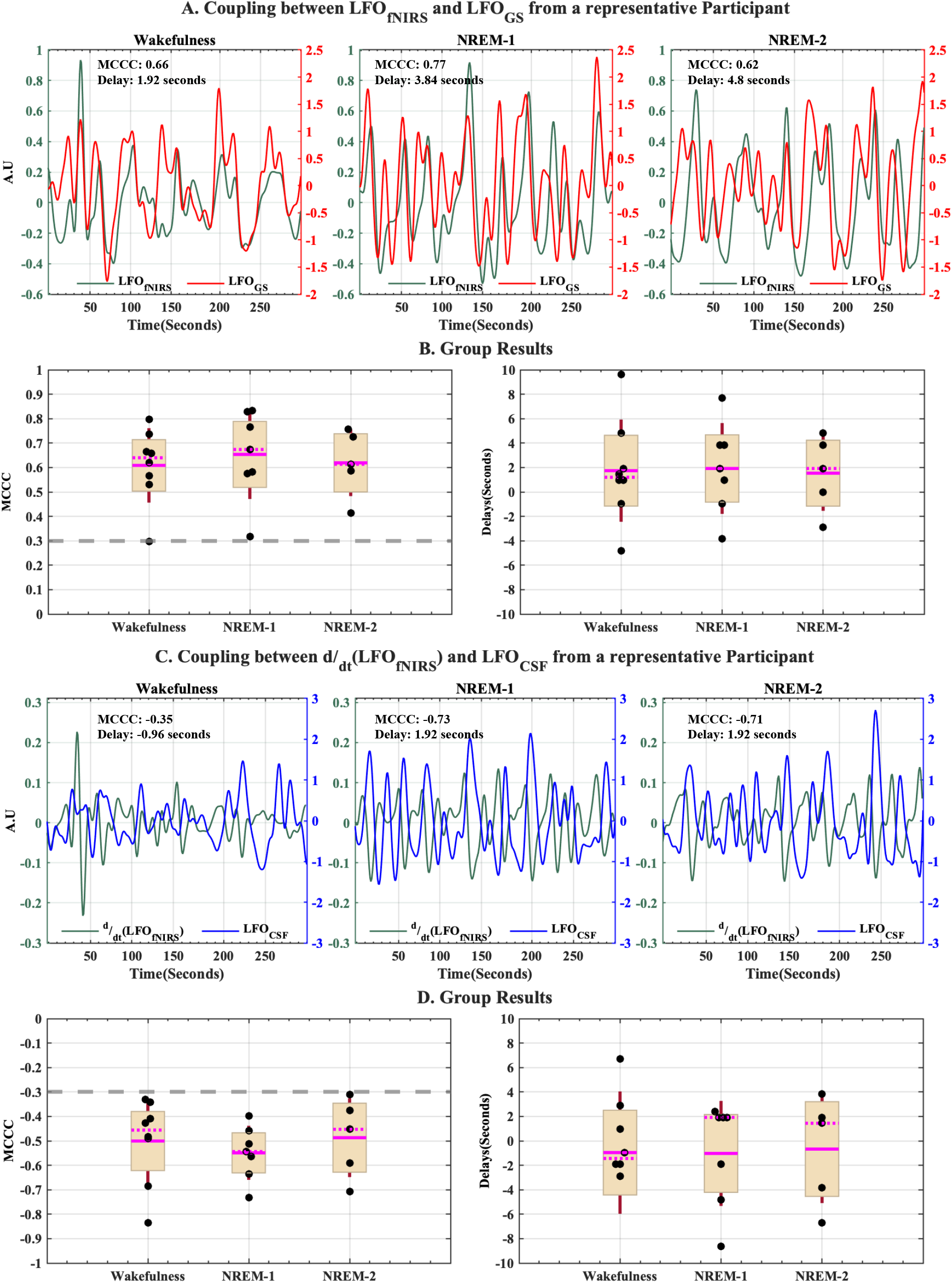
Strong coupling exists between peripheral hemodynamic LFOs and cerebral hemodynamic LFOs as well as between LF peripheral hemodynamic changes and cranially directed CSF movement across all three stages. Results of MCCCs and corresponding delays between LFO_fNIRS_ and LFO_GS_ for (A) a representative participant and (B) for all participants enrolled in the study. Results of MCCCs and corresponding delays between 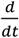 (LFO_fNIRS_) and LFO_CSF_ for (C) a representative participant and (D) for all participants enrolled in the study. A.U – Arbitrary Units; CSF – Cerebrospinal fluid; fNIRS – functional Near Infrared Spectroscopy; GS – Global Signal; LFO – Low frequency Oscillation (0.01 Hz – 0.1 Hz); MCCC – Maximum Cross-Correlation Coefficient. In group results (B) and (D), the magenta solid line represents the mean, the magenta dotted line represents the median, the brown whiskers represent one standard deviation of the raw data points jittered over a 95 percent confidence interval in cream and the gray dashed line represent the threshold of statistical significance for MCCCs in the LFO range.

Results of MCCCs and delay times between 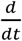 (LFO_fNIRS_) and LFO_CSF_ during continuous 5-minute segments of wakefulness, NREM-1, and NREM-2 from a representative participant are presented in Figure 6C. 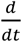 (LFO_fNIRS_) is negatively correlated (MCCC < -0.3) with LFO_CSF_ during wakefulness, NREM-1, and NREM-2. Group results (Figure 6D) confirmed this with non-significant changes in MCCCs between the three stages (wakefulness: -0.51±0.17; NREM-1: -0.55±0.11; NREM-2: -0.49±0.16) with inconsistent time delays across participants and stages.

## 4. Discussion

This study evaluated the coupling dynamics of neurofluids from resting state wakefulness through stage 2 NREM sleep – with the following significant findings: First, when compared to resting-state wakefulness, CSF movement variation only showed a relative increase during light NREM sleep. Second, a strong coupling exists between changes in cerebral hemodynamic LFOs and cranially directed CSF movement LFOs, which remained the same during wakefulness, NREM-1, and NREM-2 sleep. Third, we demonstrated that EEG SWA occurs prior to LF brain hemodynamic changes in both directions during NREM-1 and NREM-2 sleep, but not wakefulness. Finally, we demonstrated that across all stages examined, peripheral vascular LFOs from fingertips were strongly coupled with LFOs of both neurofluids. Below, we provide a physiological interpretation of these findings.

### 4.1. Neurofluid coupling is independent of the wake/sleep states

The mechanical coupling between LF changes in cerebrovascular oscillations and CSF movement at the level of the fourth ventricle in both directions (cranially directed inward/caudally directed outward) has been validated during resting state wakefulness^22,25^. Briefly, this mechanism, based on the Monro-Kellie doctrine^26^, explains that the dilation of cerebral blood vessels exerts a force on the walls of the lateral ventricle resulting in caudally directed outward CSF movement and contraction of cerebral blood vessels facilitate the cranially directed inward CSF movement, through the fourth ventricle. In this study, we show that this relationship between neurofluids in the LFO range exhibits no significant changes in coupling strength during both NREM-1 and NREM-2 (when compared to resting wakefulness). Notably, CSF movement occurs ∼2 seconds after the change in fMRI GS in all three stages. This is consistent with previous research on awake^22,25^ and asleep^12^ participants. Along with results from the current study (Figure 4), this line of work documents that the mechanical coupling between neurofluids in the LFO range in humans is independent of wake/sleep states.

### 4.2. Principal cause/pathway for neurofluid coupling in the LFO range

During resting wakefulness, several systemic physiological mechanisms, including respiration-induced natural variations in arterial CO_2_^14,15^ and spontaneous vasomotion^16,17^ contribute to cerebrovascular LFOs. At the same time, local electrocortical^12^ or autonomic neural^13^ control through neurovascular coupling have also been suggested as a possible mechanisms specific to NREM sleep. This brings us to the question: Is there a change in the principal cause behind cerebrovascular LFOs during NREM sleep compared to wakefulness? To answer this question, we explored neurofluid movement with a neurogenic origin (through the relationship between neurofluids and electrical activity of the brain) and/or a global circulatory origin (through the relationship between neurofluids and non-neuronal fingertip vascular LFOs).

#### 4.2.1. Neural origin

Fultz *et al*. reported the occurrence of EEG SWA, ∼3.8 seconds prior to fMRI blood oxygen level-dependent (BOLD) signal changes during NREM sleep (predominantly light sleep)^12^. Similar results were also illustrated during all stages of NREM sleep with differing temporal delays^13^. We extended these results and found EEG SWA peaks before thresholded 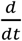 (LFO_GS_) peaks in both directions, with peak-peak delays of 2-4 seconds and 2-3 seconds respectively during NREM-1 and NREM-2 sleep (Figure 5). We also performed the same peak-locked analysis in the EEG theta (4 Hz – 7 Hz) range since it is predominant during light NREM sleep^27^ but found no active coupling (see Figure S1 in the supplementary material). Given that these coupled EEG SWA peaks were detected prior to the onset of the deep NREM-3 sleep, a potential candidate mechanism that explains our observations are autonomic regulatory pathways. The cerebral vasculature is richly innervated by both sympathetic and parasympathetic nerve fibers, and the reciprocal interaction between them ensures normal maintenance of cerebral perfusion^28^. During light NREM sleep, transient increases in sympathetic activity^29^, to mitigate the sleep-induced hypoventilation (and consequent hypercapnia)^30^, leads to cerebral vasoconstriction, as a protective measure against cerebral hyperperfusion^31^. This, in turn, leads to craniad CSF movement, per the observed coupling relationship. However, the increase in arterial blood pressure during sympathetic drive would activate the baroreceptors and lead to cerebral vasodilation via parasympathetic activation^31,32^. In turn, this vasodilation leads to caudad CSF movement. These ventilatory oscillations continue until the eupneic CO_2_ level for sleep is established^33^ and so must be the regulating autonomic pathways. This might also explain the significant increase in power of neurofluid LFOs (NREM-2 compared to wakefulness - see Figure S2 in the supplementary material) in the narrow range of 0.02 Hz – 0.04 Hz, attributed to autonomic neural origin^34^. Therefore, it appears that there is a higher autonomic neural contribution to neurofluid LFOs during light NREM sleep compared to wakefulness. However, its primary purpose is not the enhancement of CSF movement and does not lead to any significant increase in flow-weighted CSF movement fluctuations at the level of the fourth ventricle (see Figure 3).

#### 4.2.2. Systemic circulatory origin

Peripheral hemodynamic LFOs recorded (at the fingertips using fNIRS) are strongly correlated with fMRI BOLD cerebrovascular LFOs during resting state wakefulness^35,36^. In this study, we found peripheral vascular LFOs maintain a strong correlation with both neurofluid LFOs (LFO_GS_ and LFO_CSF_) during light NREM sleep as well (Figure 6). To our knowledge, this is the first study to establish this relationship during light NREM sleep. These peripheral fNIRS-based vascular LFOs represent non-neuronal systemic physiological processes with an endogenous circulatory origin at or before the heart/lung and are posited to travel with the blood^36,37^. In another study, these peripheral LFOs were shown to contain information spatially and temporally distinct from the simultaneously recorded cardiac and respiratory regressors^24^. The time delays between peripheral fNIRS LFOs and neurofluid LFOs in our study are distinct for each participant. This is consistent with previous research^35,36^ and may be explained by individual differences in length, diameter, and elasticity of blood vessels, as well as differences in height, age, and gender of the participants. Taken together, the variability in cerebral hemodynamic LFOs and the coupled CSF movement LFOs are explained by non-neuronal systemic physiology and appears to be independent of the wake/sleep states. More importantly, our data establish that the contribution of non-neuronal systemic physiology towards coupled neurofluid LFO dynamics does not change even when an alternate autonomic neural contribution comes into play during light NREM sleep. This finding has the potential to be developed as a reliable biomarker of circulatory dysfunction, thus offering the possibility of a promising non-invasive bedside tool for overnight monitoring of sleep-related vascular disorders.

### 4.3. Limitations

A limitation of this study is the lack of data on deep NREM-3 sleep and REM sleep. It would be interesting to examine how the reported neurofluid dynamics and coupling with neural/systemic ‘effectors’ change when the brain/body switch to these sleep states. Another minor limitation is the relatively small sample size and lower fMRI temporal resolution, which made it difficult to sample the cardiac pulsations in 6 participants. Future research can build on this work by considering all sleep stages.

## 5. Conclusion

In conclusion, our study documented a coupling between LF cerebral hemodynamics and CSF movement irrespective of wakefulness or light NREM sleep. It also illustrated how non-neuronal systemic physiology significantly influences neurofluid LFOs even when autonomic neural contributions come into play during light NREM sleep. A coupling index, therefore, could be developed as a reliable indicator of vascular health with applications in overnight sleep monitoring.

## Funding

This work was supported by the seed grant from Purdue Institute for Integrative Neuroscience (PI: A. J. Schwichtenberg) and National Institutes of Health grants R21 (R21AG068962, PI: Yunjie Tong) and S10 (S10 OD012336 - 3T MRI Scanner dedicated to Life Sciences Research PI: Ulrike Dydak).

## Author Contribution Statement

A. J. Schwichtenberg, Yunjie Tong, and Vidhya Vijayakrishnan Nair conceived of the presented idea. A. J. Schwichtenberg and Pearlynne L H Chong helped in participant recruitment. Vidhya Vijayakrishnan Nair, Brianna R Kish, and Ho-Ching (Shawn) Yang performed data collection. Vidhya Vijayakrishnan Nair and Brianna R Kish performed data analysis. A. J. Schwichtenberg, Yunjie Tong and Vidhya Vijayakrishnan Nair interpreted the results and developed the theory. Vidhya Vijayakrishnan Nair took the lead in writing the manuscript. Vidhya Vijayakrishnan Nair, Brianna R Kish, Pearlynne L H Chong, Ho-Ching (Shawn) Yang, Yu-Chien Wu, Yunjie Tong and, A. J. Schwichtenberg provided critical feedback and helped shape the research, analysis and manuscript.

## Disclosure/Conflict of Interest

The Author(s) declare(s) that there is no conflict of interest.

## Supplementary Material

### 1. Simulation of Motion Effects

**Table S1:** Correlations between motion parameters (FSL MCFLIRT) and cranially directed CSF signals for each participant. From the results, it can that there exist no noteworthy correlations between motion parameters and the CSF signals. This therefore confirms that the CSF signals extracted from the fMRI scans were not affected by motion. r_(x-z)_: MCFLIRT estimated rotations; t_(x-z)_: MCFLIRT estimated translation

| Parameter<br>Participant # | $r_x$ | $r_y$ | $r_z$ | $t_x$ | $t_y$ | $t_z$ |
| --- | --- | --- | --- | --- | --- | --- |
| 1 | -0.1977 | 0.1612 | -0.0496 | -0.0407 | -0.0847 | 0.1524 |
| 2 | -0.0229 | -0.0039 | -0.0332 | -0.0302 | -0.0102 | 0.0208 |
| 3 | -0.0345 | 0.0177 | 0.0071 | -0.0142 | -0.0203 | 0.0427 |
| 4 | 0.0071 | -0.0322 | -0.0076 | -0.0016 | 0.0044 | -0.0479 |
| 5 | -0.0101 | 0.0048 | 0.0130 | 0.0254 | 0.0312 | -0.0506 |
| 6 | 0.0236 | -0.0012 | -0.0158 | -0.0103 | 0.0637 | 0.0017 |
| 7 | 0.0051 | -0.0041 | -0.0236 | -0.0281 | -0.0111 | -0.0256 |
| 8 | -0.0403 | 0.0057 | 0.0026 | 0.0032 | 0.0716 | -0.1476 |
| Mean ± SD | -0.0338±0.07 | 0.0185±0.06 | -0.0135±0.03 | -0.0121±0.03 | 0.0056±0.06 | -0.0068±0.09 |

### 2. No clear theta activity peaks before low-frequency changes in fMRI global signal at any NREM sleep stage

**Figure S1:**
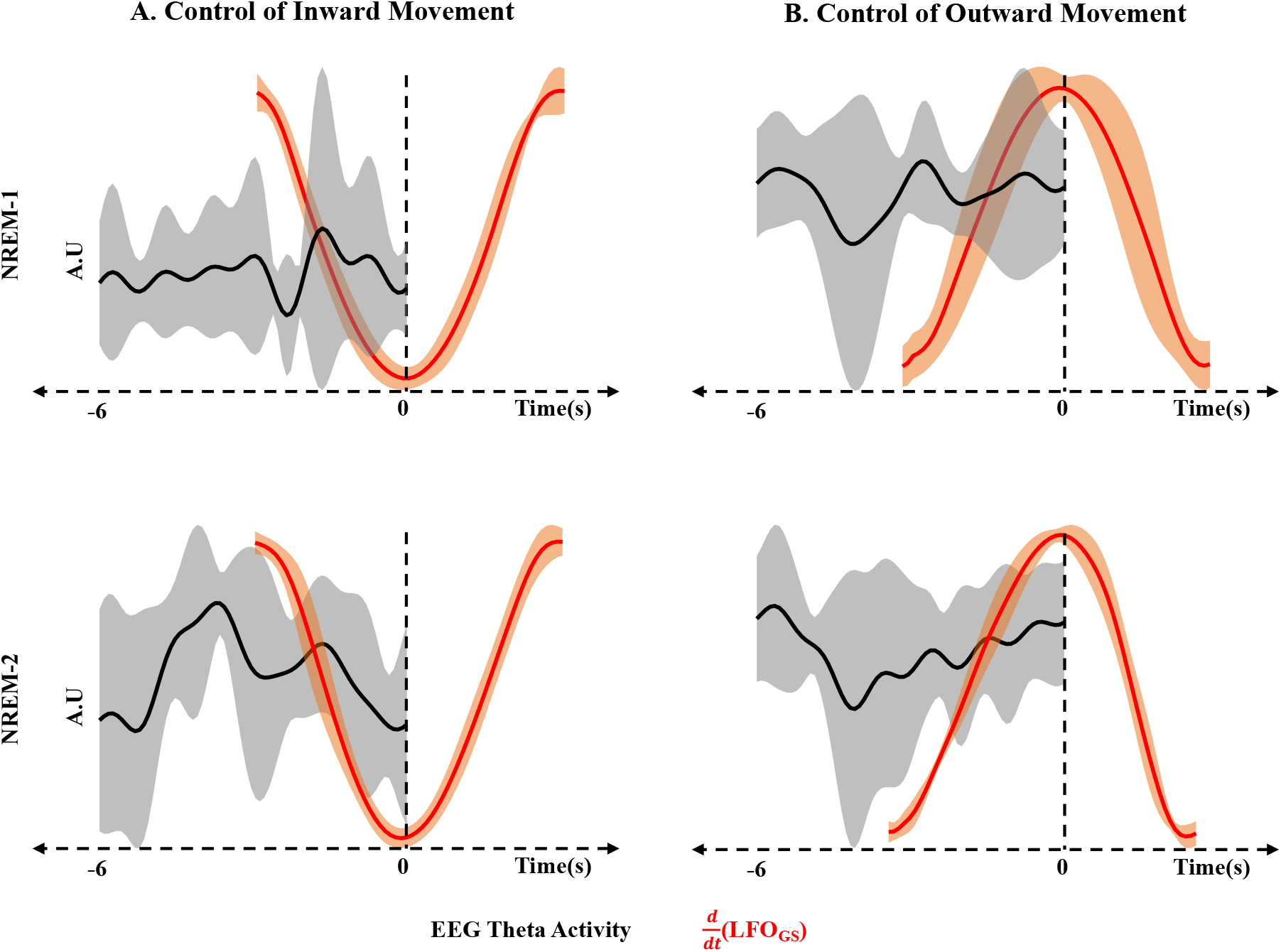
Results of 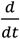 peak-locked folding average analysis of EEG theta range (4 Hz – 7 Hz) signals in (A) positive direction (cranially directed CSF inward movement) and (B) negative direction (caudally directed outward CSF movement) during N1-sleep and N2-sleep. The mean signal in each case is illustrated with thick black and red lines with standard deviation across participants represented by gray and orange regions around the mean signal. A.U – Arbitrary Units; CSF - Cerebrospinal Fluid; EEG – Electroencephalogram; GS – Global Signal; LFO – Low frequency Oscillation (0.01 Hz – 0.1 Hz).

### 3. Average power of CSF movement and GS signals in the range of 0.02 Hz – 0.04 Hz increase significantly during NREM-2 compared to wakefulness

In order to specifically understand the power variations of CSF movement and fMRI GS in the LFO range (wakefulness vs. light NREM sleep) and specifically identify the band which is responsible for any significant changes, the integrated average power estimates of these signals within the entire LFO range and specific narrow frequency ranges within the LFO band (0.01 Hz – 0.02 Hz, 0.02 Hz – 0.04 Hz and 0.04 Hz – 0.1 Hz) was quantified for every state (MATLAB bandpower). These specific frequency ranges within the LFO band were chosen based on previous research^1^ which identified them to be of endothelial, neurogenic and myogenic origin, respectively. Statistical significance in power estimates between groups were assessed using non-parametric Kruskal-Wallis test (one sample Kolmogorov-Smirnov test was used to test for normality).

The results of this power analysis of cranially directed CSF movement and fMRI GS signals during wakefulness, NREM-1 and NREM-2 are illustrated in figure S2. The individual CSF power spectrum in the LFO range of a representative participant across all the three stages (figure S2-A) shows that there is a gradual increase in the average power (integrated area under the power spectrum curve) from wakefulness to NREM-2. Group results (figure S2-B, upper panel) shows that there is a significant increase in CSF power (p-value < 0.05) in the LFO range during NREM-2 compared to wakefulness.

Further probing into specific narrow frequency bands within the LFO range (figure S2-B, lower panel) shows that this significant increase in power during NREM-2 compared to wakefulness is mostly occurring in the narrow range of 0.02 Hz – 0.04 Hz (p-value < 0.01). A similar significant increase in power in the range of 0.02 Hz – 0.04 Hz during NREM-2 can also be seen in the case of fMRI GS, although there was only a relative increase in the entire LFO range from wakefulness to N2-sleep.

**Figure S2:**
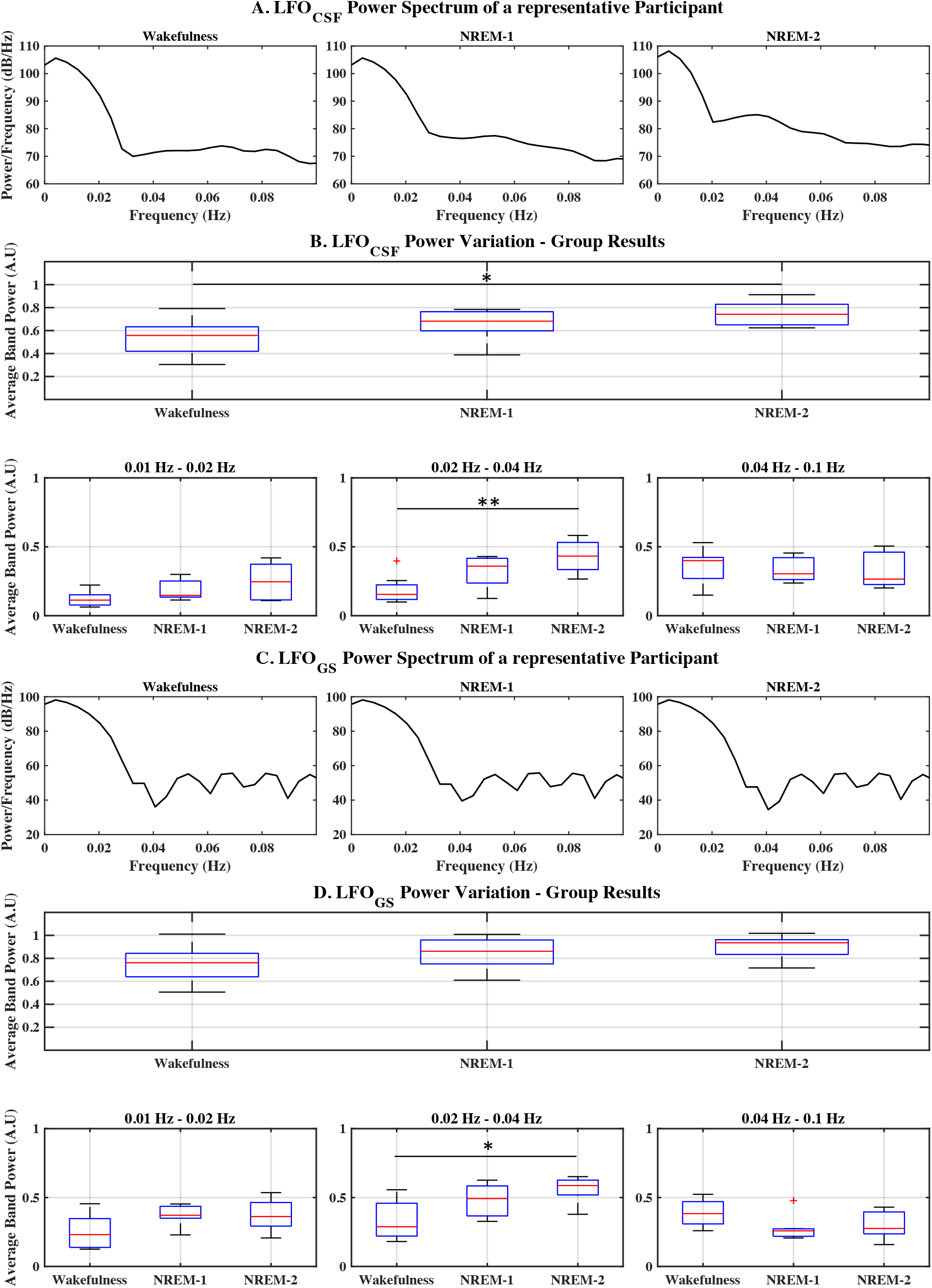
LFO range power variations in cranially directed CSF movement and fMRI global signals during wakefulness, NREM-1 and NREM-2. An example of (A) CSF power spectrum and (C) fMRI global signal in the LFO range of a representative participant and group results of (B) average LFO_CSF_ and (D) average LFO_GS_ band power from all participants enrolled in the study, across all three stages. A.U – Arbitrary Units; CSF - Cerebrospinal Fluid; GS – Global Signal; LFO – Low frequency Oscillation (0.01 Hz – 0.1 Hz); NREM – Non Rapid Eye Movement sleep; * (p-value < 0.05); ** (p-value <0.01).

## Footnotes

1 We refer to cerebral blood volume changes and CSF movement as ‘neurofluids’ in this paper, although this umbrella term may also include interstitial fluid^38^.

## Notes

### Competing Interest Statement

The authors have declared no competing interest.

